# Control of tissue development by cell cycle dependent transcriptional filtering

**DOI:** 10.1101/2020.02.25.964650

**Authors:** Maria Abou Chakra, Ruth Isserlin, Thinh Tran, Gary D. Bader

## Abstract

Cell cycle duration changes dramatically during development, starting out fast to generate cells quickly and slowing down over time as the organism matures. The cell cycle can also act as a transcriptional filter to control the expression of long genes which are partially transcribed in short cycles. Using mathematical simulations of cell proliferation, we identify an emergent property, that this filter can act as a tuning knob to control cell fate, cell diversity and the number and proportion of different cell types in a tissue. Our predictions are supported by comparison to single-cell RNA-seq data captured over embryonic development. Evolutionary genome analysis shows that fast developing organisms have a narrow genomic distribution of gene lengths while slower developers have an expanded number of long genes. Our results support the idea that cell cycle dynamics may be important across multicellular animals for controlling gene expression and cell fate.

## Introduction

A fundamental question in biology is how a single eukaryotic cell (e.g. zygote, stem cell) produces the complexity required to develop into an organism. A single cell will divide and generate many progeny, diversifying in a controlled and timely manner (Mueller et al., 2015) to generate cells with very different functions than the parent, all with the same genome (Wilmut et al., 1997). Many regulatory mechanisms coordinate this process, but much remains to be discovered about how it works (Zoller et al., 2018). Here, we explore how cell cycle regulation can control gene expression timing and cell fate during tissue development.

The canonical view of the cell cycle is a timely stepwise process. Typically, the complete cell cycle is divided into four phases: first gap phase (G1), synthesis phase (S), second gap phase (G2) and mitotic phase (M). The length of each phase determines how much time a cell allocates for processes associated with growth and division. However, the amount of time that is spent in each phase frequently differs from one cell type to another within the same organism. For example, some cells experience fast cell cycles, especially in early embryogenesis. Organisms such as the fruit fly *(Drosophila melanogaster*) and the worm (*Caenorhabditis elegans*) exhibit cell cycle durations as short as 8-10 mins (Edgar et al., 1994; Foe, 1989). Cell cycle duration also changes over development. For example, it increases in mouse (*Mus musculus*) brain development from an average of 8 hrs at embryonic day 11 (E11) to an average of 18hrs by E17 (Edgar et al., 1994; Foe, 1989). Duration can also differ spatially among cells in a tissue; for instance, in the fruit fly embryo, by cycle 14, cells are grouped spatially by their cell cycle duration – ranging from 8 minutes in the anterior region to 170 minutes in the mid-posterior region (Foe, 1989).

Interestingly, cell cycle duration can act as a transcriptional filter that constrains transcription (Shermoen and O’Farrell, 1991). For instance, if the cell cycle progresses relatively fast, transcription of long genes will be interrupted. In typical cells, the gene transcription rate is between 1.4-3.6 kb per minute (Ardehali and Lis, 2009). Thus, an 8 minute cell cycle would only allow transcription of the shortest genes, on the order of 10 kb measured by genomic length, including introns and exons, whereas a 10 hour cell cycle would allow transcription of genes as long as a megabase on the genome.

Cell cycle dependent transcriptional filtering has been proposed to be important in cell fate control (Bryant and Gardiner, 2016; Swinburne and Silver, 2008). Most multicellular eukaryotic animals start embryogenesis with short cell cycle durations and a limited transcription state (O’Farrell et al., 2004) with typically short zygotic transcripts (Heyn et al., 2014). These cells allocate the majority of their cycle time to S-phase (synthesis), where transcription is inhibited (Newport and Kirschner, 1982), and M-phase (division), with little to no time for transcription in the gap phases. However, as the cell cycle slows down, time available for transcription increases (Newport and Kirschner, 1982a), enabling longer genes to be transcribed (Shermoen and O’Farrell, 1991; Yuan et al., 2016).

Through extensive mathematical simulations of cell proliferation, we discover the novel and unexpected finding that a cell cycle dependent transcriptional filter directly influences the generation of cell diversity and it can provide fine-grained control of cell numbers and cell type ratios in a developing tissue. Our computational model operates at single-cell resolution, enabling direct comparison to recent single-cell RNA-seq data captured over development demonstrating that these experimental data match our predictions. Our work has the potential to shift the typical view of the cell cycle as a housekeeping process only used to create new cells to one where cell cycle parameters are important regulators of development.

## Results

### Computational model of multicellular development

We model multicellular development starting from a single totipotent cell that gives rise to many progeny, each with different transcriptomes that can define their cell type or state. We developed a single cell resolution agent-based computational model to simulate this process (see materials and methods). Each cell behaves according to a set of rules, and cells are influenced solely by intrinsic factors (e.g. number of genes in the genome, gene length, transcript levels and transcription rate). We intentionally start with a simple set of rules, adding more rules as needed to test specific mechanisms. We omit external cues (e.g. intercellular signaling or environmental gradients) to focus on the effects of intrinsic factors.

In our model, each cell is characterised by a fixed genome containing a set of G genes (gene_1_, gene_2_, …. gene_G_), shown in Figure 1A. Each gene_i_ is defined by a length, L_i_ (in kb), and in all our simulations each gene is assigned a different length (L_1_ < L_2_ <…L_G_). Since each gene_1_ has a unique length, L_i_, we label genes by their length (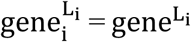; e.g. gene^3^ is a gene of length 3 kb). We assume transcription time for gene_i_ is directly proportional to its length, L_i_. In the model, each cell_j_ is initialized with a cell cycle duration (Γ_cell_) which represents the total time available for gene transcription. For example, we can initialise cell_1_ with a three-gene genome (gene^1^, gene^2^, gene^3^), where L = (1 kb, 2 kb, 3 kb) and a cell cycle duration Γ_1_ of 1 hr. We fix transcription rate, λ, to 1 kb/hr for all genes. As transcription progresses for all genes, cell_1_ will only express gene^1^. Increasing cell cycle duration, Γcell, will allocate more time for transcription, allowing longer genes to be transcribed. For example, if we initialise cell_2_ with a cell cycle duration Γ_2_=3 hrs, cell_2_ will express all three genes, with time to make three copies of gene^1^ (Figure 1B).

**Figure 1:**
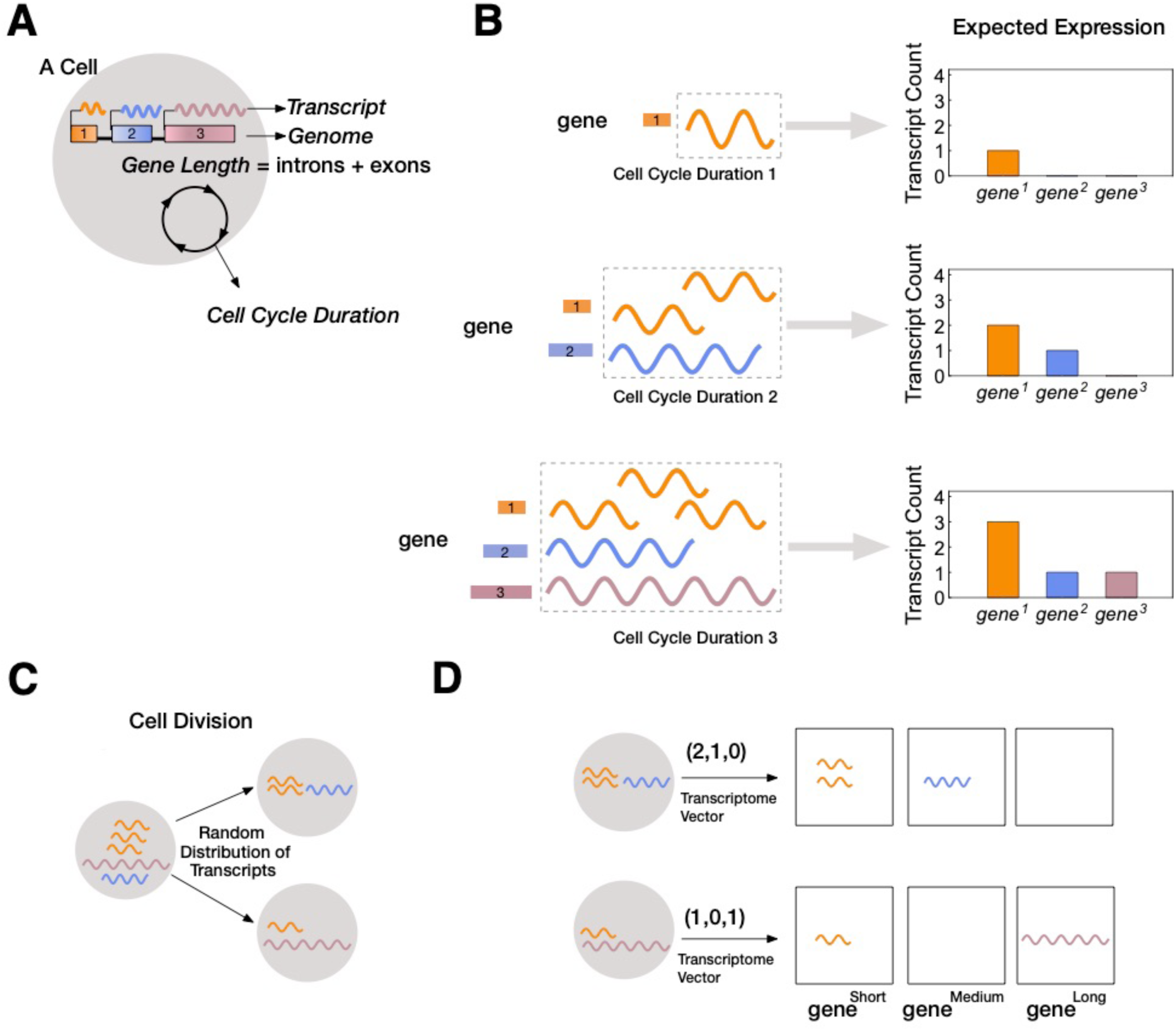
A) An illustration of a cell with three genes in its genome and gene lengths gene^1^ < gene^2^ < gene^3^. Cell cycle duration defines the time a cell has available to transcribe a gene. B) A cell with cell cycle duration = 1 hr will only enable transcription of gene^1^; cell cycle duration = 2 hrs will enable transcription of gene^1^ and gene^2^; cell cycle duration = 3 hrs enables transcription of all three genes. C) Our model assumes transcripts passed from parental cell to its progeny will be randomly distributed during division (M phase). D) Each cell is characterized by its transcriptome.

Once transcription is complete, the cell enters M-phase, during which it divides, and expressed transcripts are randomly distributed to the two progeny cells (Figure 1C). We assume that transcription begins anew at the start of the cell cycle (i.e. all transcripts from a gene that can’t be finished are eliminated), modeling the known degradation of incomplete nascent transcripts in M-phase (Shermoen and O’Farrell, 1991). We can relax our assumption to include parental transcript inheritance and decay where a proportion of inherited parental transcripts are removed after each cell division (Sharova et al., 2009). All cells and their transcriptomes are tracked over the course of the simulation for analysis, enabling single cell resolution analysis. Transcriptomes are stored as vectors containing the total number of transcripts per gene. For instance, cell_2_ may have a transcriptome of (3,1,1), indicating that three genes are expressed, with gene^1^ expressed at three transcripts per cell and the other two genes expressed at one transcript per cell (Figure 1D).

### Cell cycle duration influences transcript count: short genes generate more transcripts than longer genes

We begin by examining how a transcriptional filter impacts transcript counts (gene expression levels), as controlled by cell cycle duration. Shorter cell cycles will interrupt long gene transcription resulting in relatively high expression of short genes and low expression of long genes. Our computational simulations recapitulate this expected pattern (Figure 2A). Each simulated cell transcriptome is divided into three bins containing short, medium and long genes and then each bin is summarized with an average transcript count. In simulations, bins with short genes exhibit the highest expression levels. As cell cycle duration increases, more cells show an increase in expression of longer genes (Figure 2A and S1).

**Figure 2:**
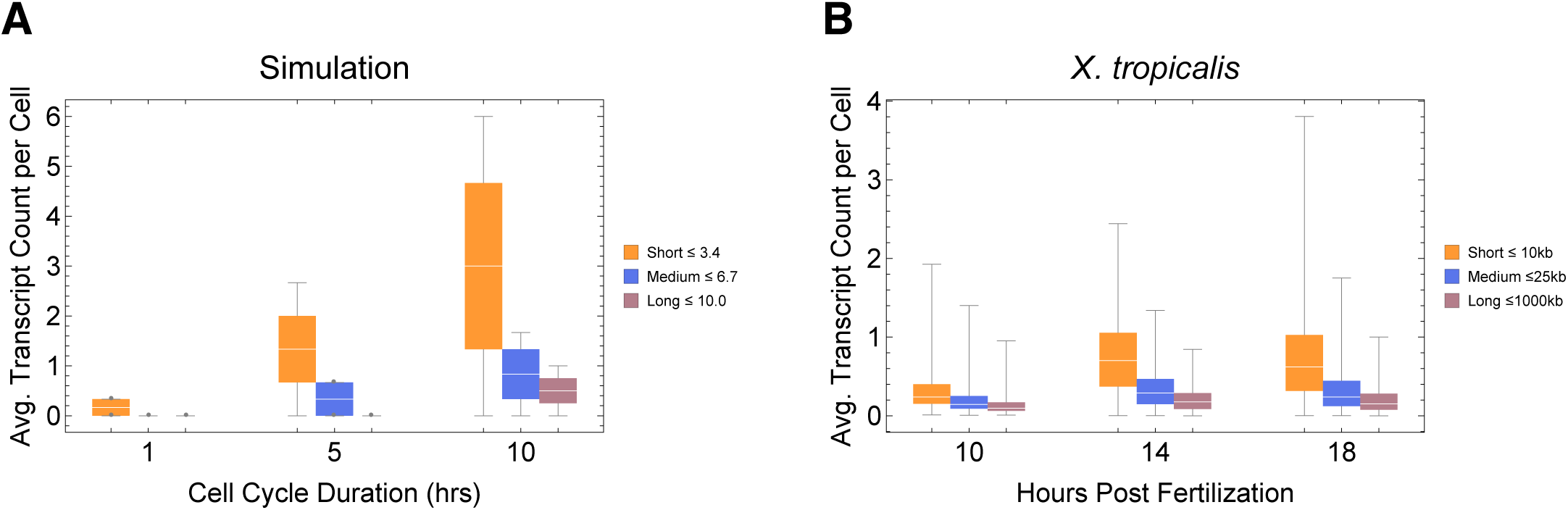
Short genes produce more transcripts than longer genes at multiple cell cycle duration lengths. The transcriptome for each cell is subdivided into short, medium, and long gene bins and transcript counts are averaged per bin. A) Simulations predict that short genes will be more highly expressed than long genes. Simulation results are shown for cell cycle durations of 1, 5 and 10 hrs (other parameters genome G = 10 (gene^L^_1_^-L^_G_) L_1_=1, L_G_=10, ploidy=1, one cell division, iterations = 20000). B) *Xenopus tropicalis* single cell data obtained from GSE113074 displaying expected patterns where short genes (lengths <10 kb, ∼4174 genes) have a higher expression than both medium genes (lengths > 10kb, ∼3607 genes) and longer genes (lengths >25 kb, ∼3049 genes).

Single cell RNA-seq (scRNA-seq) has recently been used to profile gene expression of thousands of cells over multiple embryonic developmental time points in *Xenopus tropicalis* (Briggs et al., 2018) and *Danio rerio* (Wagner et al., 2018). These systems have relatively short cell cycle durations achieving developmental hatching time as short as 6 and 3 days, respectively (Kimmel et al., 1995). We analyzed these data in the same manner as our model (Figure 2B, and S2) and found that, in general, cells express more short genes than longer genes over multiple developmental time points. Thus, gene expression patterns from multiple scRNA-seq developmental time course data sets are compatible with our model prediction.

### Cell cycle duration can control cell diversity

We next asked how three major model parameters (cell cycle duration, maximum gene length, and number of genes in the genome) can influence the generation and control of cell diversity observed during normal multicellular development. We conducted simulations for a single cell division step for simplicity, but these were repeated thousands of times to model cell population effects. We compute cell diversity in two ways; first, as the number of distinct transcriptomes in the cell population (transcriptome diversity); and second, as the number of distinct transcriptomic clusters, as defined using standard single cell transcriptomic analysis techniques (Satija et al., 2015). Both measures model real cell types and states that are distinguished by their transcriptomes, with transcriptome diversity as an upper bound on cell type number and cluster number as a much reduced lower bound. We first ran simulations with an active transcriptional filter by varying only the cell cycle duration, Γ, for a genome with 10 genes, ranging in size from 1 to 10 kb such that it satisfies L_1_ = 1 ≤ … Γ… ≤ L_G_ = 10. Short cell cycle duration parameter values generated a homogenous population of cells because only short transcripts can be transcribed. As cell cycle duration was increased, transcriptome diversity also increased. Longer cell cycle duration values generated heterogeneous populations, because a range of transcripts can be expressed (Figure 3A, brown line). Interestingly, cell cluster diversity peaks at intermediate cell cycle duration parameter values (Figure 3B, brown line; 3C), because new genes are introduced with increasing cell cycle lengths, but eventually long cell cycles provide sufficient time for cells to make all transcripts which leads to reduced variance between the progeny. We next repeat this experiment by turning off the transcriptional filter by reducing the maximum gene length such that L_G_ < Γ, (Figure 3A,B, blue line). In this case, cell diversity can be generated, but it quickly saturates, as all genes are expressed and duplicated with additional 1 hour increases in cycle duration. Thus, while cellular diversity can be generated with an active or inactive transcriptional filter, only when the transcriptional filter is active can diversity be controlled by cell cycle duration.

**Figure 3:**
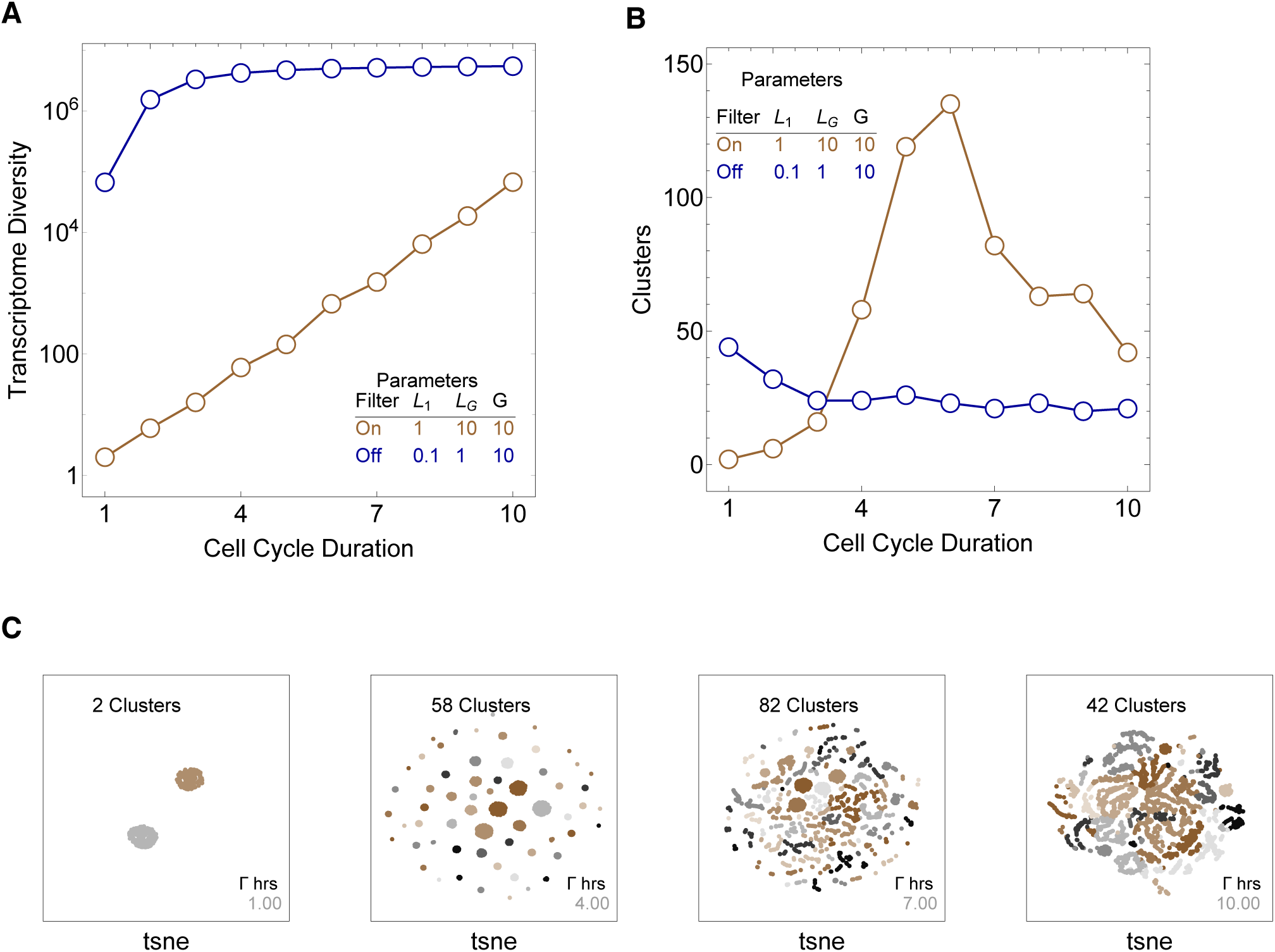
Simulations exploring effects of cell cycle duration, Γ, gene number, G, and gene length distribution. A) Simulations show that cell diversity (transcriptome diversity) increases as a function of cell cycle duration. Short cell cycle durations can constrain the effects of gene number as long as a transcriptional filter is active (gene length distributions are broad, L_1_ < … (Γ ∗ λ) … < L_G_). When L_G_ < (Γ ∗ λ), cell cycle duration does not control cell diversity. Cell cycle duration effects are relative to the distribution of gene length in the genome. B) We use Seurat to cluster the simulated single cell transcriptomes and report the number of cell clusters over the simulations. This shows that cell diversity increases with gene number but the number of clusters identified decreases when all the genes can be expressed similarly among all cells. C) Representative examples of t-SNE visualizations are shown for simulations with cell cycle durations 1, 4, 7 and 10 hrs (genome G = 10, gene^L^_1_^-L^_G_, ploidy n = 1 and transcription rate, λ = 1 kb/hr).

We next performed extensive simulations with various parameters to generalize our understanding of how cell diversity can be created (Table S1). The results enabled us to determine an analytical solution of how cell transcriptome diversity can be generated in our model that accurately predicts simulation results (Table S1). Cell diversity increases as a function of cell cycle duration (Γ), transcription rate (λ), and number of genes in the genome (G). In particular, 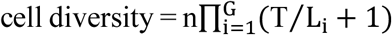, where n is the genome ploidy level, T = Γ ∗ λ (i.e. the maximum transcribed gene length, T, is restricted by the product of cell cycle duration, Γ, and transcription rate, λ), and L_i_ is the length of gene_i_. While the number of genes and their length distribution can change over the course of evolution, these numbers are constant for a given species, and transcription rate is likely highly constrained (Ardehali and Lis, 2009), leaving only cell cycle duration as a controllable parameter of cell diversity during development according to our model.

### Gene length distribution of a genome correlates with cell cycle duration during development

To examine whether gene length distributions in real genomes are compatible with our transcriptional filter model, we examined how these distributions across a variety of genomes relate to cell cycle timing during development. Our model predicts that if an organism has longer genes, it must have longer cell cycles during development to accommodate them. Unfortunately, measurements of cell cycle timing across species are difficult to find, however cell cycle duration should be broadly correlated with overall development time, which is readily available. In particular, we expect faster developing organisms to have shorter cell cycles and genes whereas slower developing organisms will have longer cell cycles and genes. We analyze gene length distributions for 11 genomes spanning budding yeast to human with a diverse range of developmental durations as shown in Figure 4 and S3, Table S2 (Gilbert and Barresi, 2016; Jukam et al., 2017). Non-mammalian species that we analyze are relatively fast developing, ranging from approximately two hours (e.g. *S. cerevisiae*) to a few days (e.g. *X. tropicalis* and *D. rerio*), while mammals (*Mus musculus, Sus scrofa, Macaca mulatta*, and *Homo sapiens*) are relatively slow developers (20, 114, 168 and 280 days, respectively). In agreement with our model, the gene length distribution is narrower and left shifted (shorter genes) for fast developers and broader and right shifted (longer genes) for slower developing species. To illustrate this quantitatively, using a typical transcription rate of 1.5 kb/min (Ardehali and Lis, 2009), a cell cycle duration of 1 hr can exclude 14% of the total genes found in relatively slow developers and 0.8% of relatively fast developers (Figure 4). These results support the idea that cell cycle duration is important for controlling gene expression during development *in vivo* and this effect could be generally important across multicellular animals.

**Figure 4:**
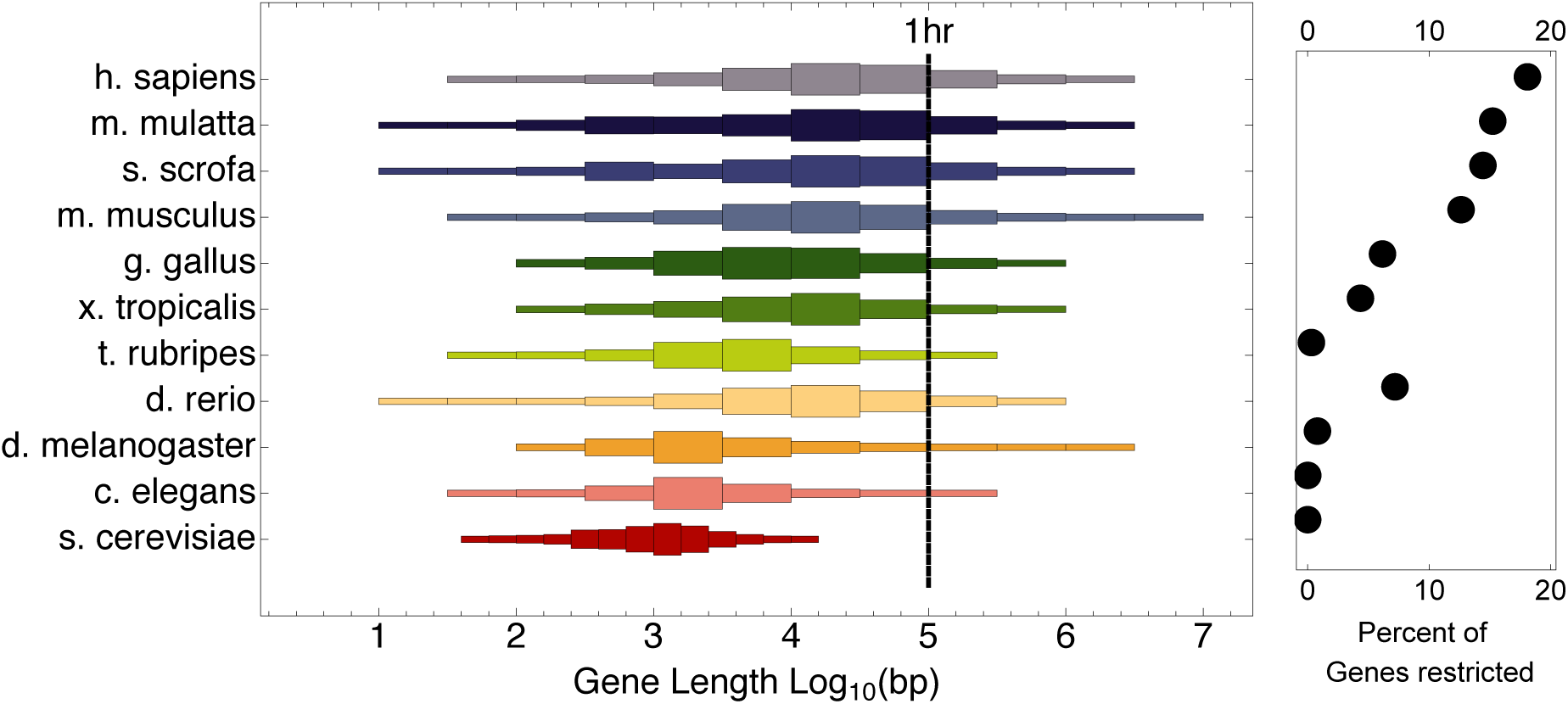
Model organisms exhibit a large diversity in gene length distributions over their genomes. Species that have narrower gene length distributions tend to develop faster, while slow developers (mammals) exhibit broad and right shifted gene length distributions. Demarcating a 1 hr cell cycle duration using an average transcription rate of 1.5 kb/min illustrates the proportion of genes that would be interrupted before completion for each organism. *Saccharomyces cerevisiae* (budding yeast), *Caenorhabditis elegans* (worm), *Drosophila melanogaster* (fruit fly), *Danio rerio* (zebrafish), *Takifugu rubripes* (fugu), *Xenopus tropicalis* (frog), *Gallus gallus* (chicken), *Mus musculus* (mouse), *Sus scrofa* (pig), *Macaca mulatta* (monkey), *Homo sapiens* (human).

### Changing cell cycle duration over developmental time controls tissue cell proportions and number

During multicellular organism development, it is essential to generate the correct numbers of cells and cell types (diversity). Cell cycle duration changes dramatically during development, generally starting out fast to generate cells quickly and slowing down over time as the organism matures (Farrell and O’Farrell, 2014; O’Farrell et al., 2004). Clearly, cells with short cell cycles generate more progeny compared to those with longer cell cycles. However, we propose that a trade-off exists, balancing the generation of diversity (longer cell cycle durations) with the fast generation of cells (shorter cell cycle durations) (Figure 3B). To study this trade-off, we simulated cell propagation under a scenario where, after the first division, one child cell and its progeny maintains a constant cell cycle duration (Γ_1_ = 1 hr) and the second child cell and its progeny maintains an equal or longer cell cycle duration (six simulations were run for a range of cell cycle durations of the second cell, Γ_2_, from 1 to 4 hrs) over 100 total cell divisions. We initialize the starting cell with no prior transcripts (naïve theoretical state) and a genome containing five genes ranging from length one to two kb (gene^1^, gene^1.25^, gene^1.5^, gene^1.75^, gene^2^), enabling an active transcriptional filter when cell cycle duration is between one and two, matching the smallest and largest genes.

In the simulation where both cell lineages cycle at the same rate, both lineages generate the same number of progeny with the same level of diversity (Figure 5A, leftmost data point). As cell cycle duration for the second lineage and its progeny (blue line) is increased across simulations, the transcriptional filter acts to generate more diverse progeny (blue line peak), until the cell cycle is long enough to remove the transcriptional filter and cell cluster diversity is reduced (Figure 5A, blue line). Meanwhile, the short cell cycle lineage maintains a steady, low level of diversity generation (Figure 5A, gray line). Examining the generated single cell transcriptomes using clustering and t-SNE plots (Figure 5B) shows that the short cell cycle duration lineage (gray cells) only contributes to two clusters (naïve cell with transcriptome vector = (0,0,0,0,0) and cells expressing the shortest gene with transcriptome vector = (1,0,0,0,0)). The longer cell cycle duration lineage (blue cells) contributes to an increasing number of clusters over the transcriptional filter range, but with more limited cell numbers and progressively smaller population proportions (pie charts) due to the slower cell cycle (Figure 5B). This shows how cell cycle dependent transcriptional filtering can control a trade-off between cell number and diversity generation and, when mixed, can control both diversity and cell number.

**Figure 5:**
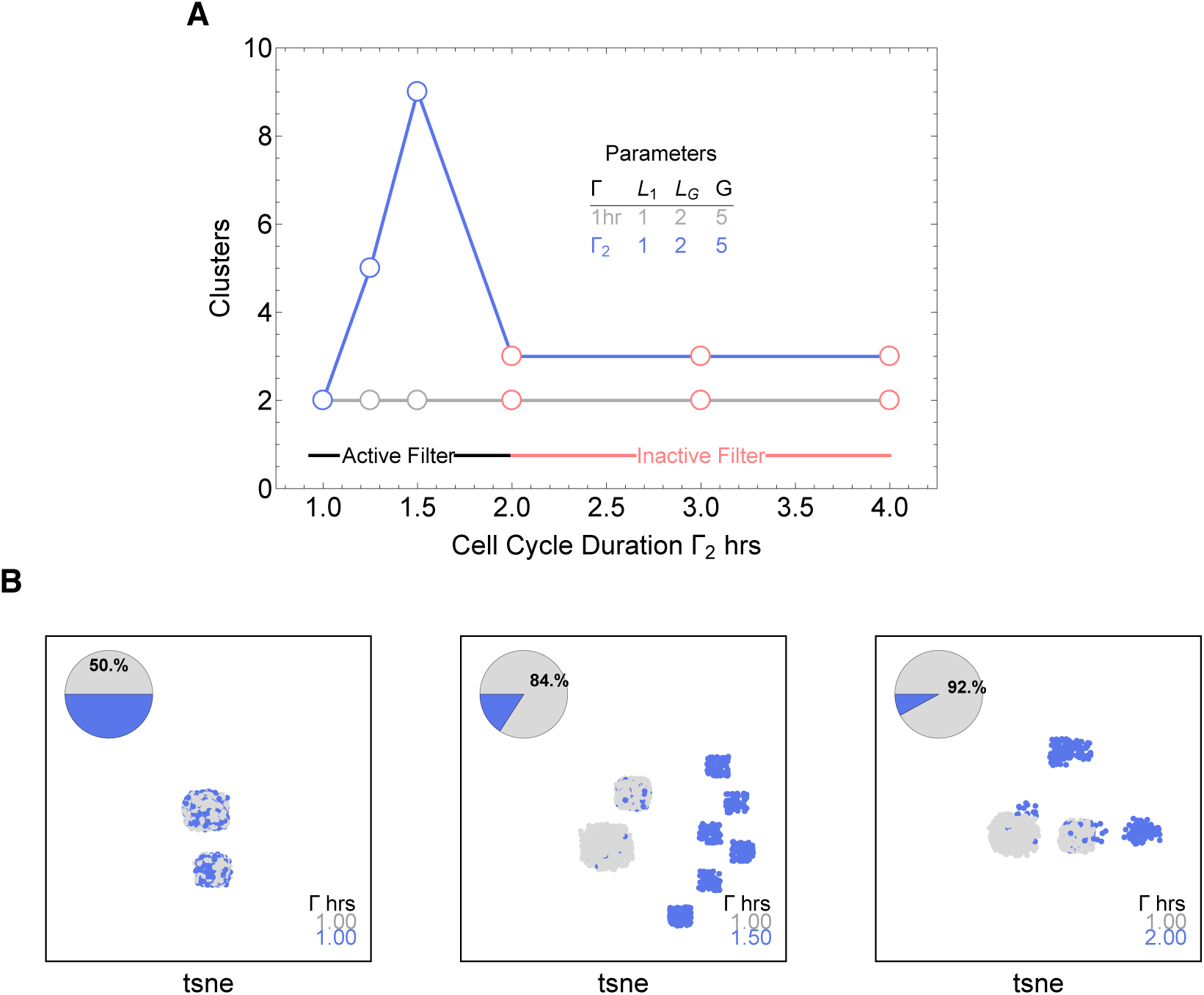
Simulations exploring the effects of cell cycle duration on number of cells and cell types. Simulations start with two cells and 100 total cell divisions are simulated (10000 simulations). Cell_1_ is initialized with cell cycle duration Γ_1_ = 1 hr, Cell_2_ has cell cycle duration, Γ_2_, ranging from 1 to 4 hrs. All progeny are tracked based on their cell cycle duration (short cell cycle, gray (Γ_1_), or longer cell cycle, blue (Γ_2_)). A) Seurat clustering shows that longer cell cycle durations for Cell_2_ increases the diversity of cells in the population (8000 randomly selected cells from the total simulation were clustered). B) Pie Chart and t-SNE visualization shows that when the cell cycle duration is the same, both cells contribute the same number of progeny and cell proportions are 50:50 (left panel). Cells with longer cell cycle duration (blue cells) generate fewer progeny with respect to the cells with a short cell cycle duration of 1 hr (gray). However the slower cells contribute more to the diversity observed in the population, shown as the blue clusters. Thus, increasing cell cycle duration increases cell diversity, but also limits the number of progeny generated.

To more faithfully simulate multicellular animal development where cell cycle duration is known to change over time, we next allowed progeny cells to differ in their cell cycle duration from their parents. We simulated increasing and decreasing cell cycle durations at various rates over up to 25 cell divisions. Increasing the cell cycle duration over time can result in a gradual increase in cell diversity (Figure 6A). The opposite trend results when cell cycle duration slowly decreases (Figure 6B). In both cases cell cycle duration dynamics affect the frequency of cells over time. Thus, cell cycle duration dynamics can exert fine-grained control of cell diversity and numbers over developmental time.

**Figure 6:**
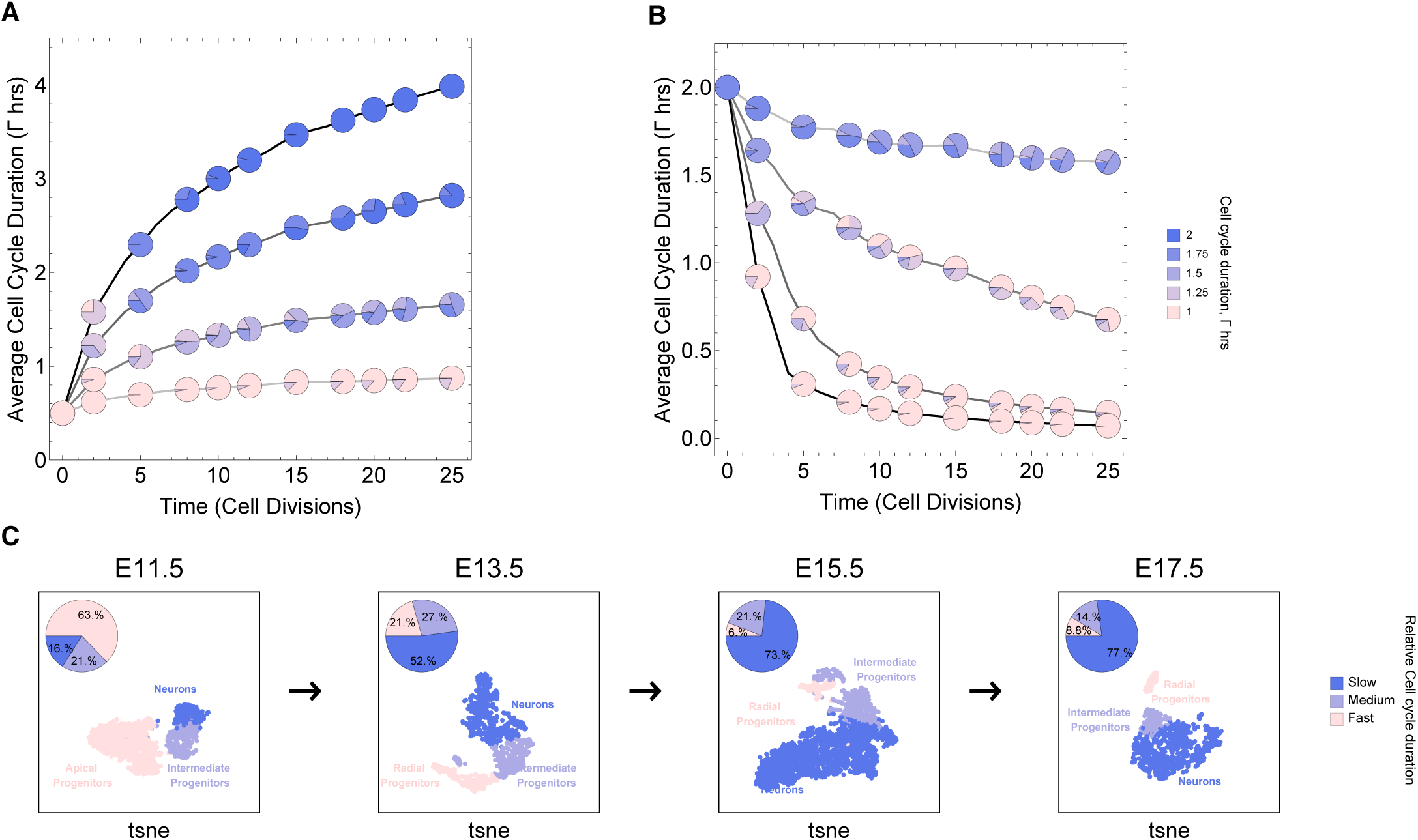
Simulations exploring the effects of cell cycle duration changes across time. All cell progeny are labeled based on their cell cycle duration (inherited from parent). Simulation of gradually A) increasing and B) decreasing cell cycle duration over time. Each line represents a different rate of increase or decrease in average cell cycle duration, following rate increments 1 (shallowest curve), 3, 6 and 9 (steepest curve). Gradual cell cycle duration changes affect the relative proportion of cells with different cell cycle durations (pie charts). Average cell cycle duration is shown in hours. Parameters Γ=Gaussian(mean Γparent ± rate increment, standard deviation σ=0.06), rate increment = 1, 3, 6 or 9, genome=(gene^1^, gene^1.25^, gene^1.5^, gene^1.75^, gene^2^), λ=1 kb/hr, cells underwent 25 divisions, iterations= 5000, ploidy n=1. C) Single cell transcriptomics data from GSE107122 (Yuzwa et al., 2017) for embryonic mouse cortex development, known to exhibit increasing cell cycle duration over time. At E11.5 the average cell cycle duration is 8 hrs and by E17.5 it is 18 hrs (Furutachi et al., 2015; Takahashi et al., 1995). Here we show how cell proportions (pie charts) change across time, with apical progenitors (relatively fast cycling cells) decreasing in frequency as the average cell cycle duration increases.

To compare with a real system, we explore single cell transcriptomics data measured over mouse cortex development (Yuzwa et al., 2017). Average cell cycle duration over mouse cortex development is known to increase from 8 hrs at embryonic day 11 (E11) to an average of 18hrs by E17 (Furutachi et al., 2015; Takahashi et al., 1995). Within this range, progenitor cells are, in general, expected to be characterized by fast cycles with short G1 duration (Vallier, 2015) and neurons by slower cell cycles with long G1 duration (Calegari et al., 2005). In the mouse cortex data, we observe an overall pattern of increasing number of cells with long cell cycle duration and decrease in fast cycling cells (Figure 6C) similar to those observed in our simulations (Figure 6A), supporting the idea that cell cycle duration dynamics could play an important role in controlling neural development.

## Discussion

Differentiation is achieved when a cell acquires a specific gene expression pattern. Many factors affect cell fate during multicellular organism development, such as cytoplasmic molecules and gradients, cell-cell communication and microenvironment signals (Edgar et al., 1986; Tabansky et al., 2013; Telley et al., 2019; Yoon et al., 2017). However, it is still unclear how cell autonomous processes support the carefully orchestrated timing of tissue development that results in a viable multicellular organism. One hypothesis is that genomic gene length can be used as a mechanism to control transcription time (Artieri and Fraser, 2014; Gubb, 1986; Keane and Seoighe, 2016; Swinburne et al., 2008). Bryant and Gardiner further propose that cell cycle duration may play a role in filtering genes that influence pattern formation and regeneration (Bryant and Gardiner, 2018; Ohsugi et al., 1997).

We explore the effect of a cell cycle dependent transcriptional filter model in development and discover the novel emergent property that the filter can be used to simultaneously exert fine grained control of the generation of cell diversity, the overall cell growth rate and cellular proportions during development. Genomic information (gene number and gene length distribution) and cell cycle duration are critical parameters in this model. Across evolutionary time scales, cell diversity can be achieved by altering gene length (Keane and Seoighe, 2016), however, in terms of developmental time scales, we propose that cell cycle duration is a major factor that controls cell diversity and proportions.

We predict that increasing the gene length distribution across a genome over evolution can provide more cell cycle dependent transcriptional control in a developing system, leading to increased cellular diversity. Examining a range of genomes and associated data supports this idea. We observe that fast developing organisms have shorter median gene lengths relative to the broad distributions including many long genes exhibited by slow developers (mammals). This aspect of genome structure may explain the observed rates of cell diversity and organism complexity, as measured by number of different cell types, over a wide range of species, Figure S3 (Valentine et al., 1994; Vogel and Chothia, 2006).

A clear pattern in real single cell transcriptomics data shows that short genes are expressed more highly than long genes, as our model predicts. However, some long genes are expressed early in development. These may be explained by inheritance from the parent cells during the maternal to zygotic transition. Indeed, if we add transcript inheritance to our model, we see the same pattern of a small number of long transcripts present early (Figure S4). The timing of zygotic transcription varies in multicellular eukaryotes (Jukam et al., 2017). Our model suggests that maternal contributions act to offset the effects of fast early cell cycles and the delayed onset of transcription. Specifically, we expect longer transcripts to be major contributors during the early maternal phase, which agrees with zebrafish (*D. rerio*) experiments showing that maternal transcripts are longer and have evolutionary conserved functions (Heyn et al., 2014).

Our analysis raises interesting directions for future work. We focus on development, but transcriptional filtering may be important in any process involving cell cycle dynamics, such as regeneration (Bryant and Gardiner, 2018), immune activation and cancer. We must also more carefully consider cell cycle phase, as transcription mainly occurs in the gap phases (Bertoli et al., 2013; Newport and Kirschner, 1982b). Experiments indicate that a cell will have different fates depending on its phase (Dalton, 2013; Pauklin and Vallier, 2013; Vallier, 2015). In our model, a cell at the start of its cell cycle will have a different transcriptome in comparison to the end of the cell cycle. Also cells with synchronous cell cycles will exhibit less variability from cells out of sync. Induced pluripotent cell state is also associated with cell cycle phases (Dalton, 2015) and efficient reprogramming is only seen in cell subsets with fast cell cycles (Guo et al., 2014). Our model could explain these observations, as slower cycling cells could express long genes that push a cell to differentiate rather than reprogram. Experimental data about cell phase measured in developing systems will help explore these effects. Further, it will be important to explore how cell cycle duration is controlled. Molecular mechanisms of cell cycle and cell size (Liu et al., 2018) control could be added to our model to provide a more biochemically realistic perspective on this topic. Ultimately, a better appreciation of the effects of cell cycle dynamics will help improve our understanding of a cell’s decision making process during differentiation, and may prove useful for the advancement of tools to control development, regeneration and cancer.

## Materials and Methods

### Mathematical model

A single cell is modeled as a sphere with a center (x,y,z) and a radius r. Alone, a single cell assumes a spherical shape, however we assume a cluster of cells will intercalate to minimise the interfacial surface area, acting like soap bubbles (Thompson, 1942). We use an established geometrical framework to calculate the distance between any number of cell centres (Isenberg, 1992). For instance, under these conditions, two cells, Cell_i_ and Cell_j_ will be 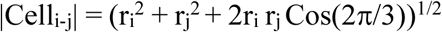 apart. The equation uses the Cosine Law to calculate the distance between centres for Cell_i_ and Cell_j_, given that their radii, r_i_ and r_j_, meet at an angle equal to π/3 radians (Isenberg, 1992).

Each cell can divide and make two progeny cells. Cells behave and interact according to a fixed set of rules. Our major rule is a gene-length mechanism, where each cell is defined by a genome and a cell cycle duration. The cell cycle duration determines which genes are expressed within the cell. All decisions are based on a cell’s autonomous information and we omit external factors. We deliberately choose to consider a simple baseline setup (without additional mechanisms) in which the effect of cell cycle duration becomes most transparent.

Each cell is defined by position (x,y,z), a genome size G, cell cycle duration in hours and the transcripts inherited or recently transcribed. In the genome, each gene is defined by a length, gene^Length^. For example of a genome with three genes (gene^1^, gene^2^, gene^3^) represents genes of length 1, 2, 3 kb, respectively.

One cell division (one time step): Each Cell_i_ will transcribe its genes based on the time available, defined by the cell cycle duration. Once a cell cycle is finished, the cell divides. When cells are synchronized, the first cell division T= Γ_i_. When the cells are asynchronized then the algorithm identifies the time allocated as the shortest cell cycle duration in the population. The number of transcripts for each gene is calculated by a function of cell cycle duration, Γ, transcription rate, λ and gene length: 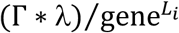. If the cell does not divide, then the current cell cycle phase is computed and stored. If the cell can divide within the time T then it will randomly, using a uniform distribution, assign its transcripts amongst its two progeny cells. For certain experiments, the cell cycle duration for each progeny is allowed to diverge from the parental duration using a monotonic function (increasing or decreasing) and a stochastic variable based on a Gaussian distribution with a mean equal to Γ_i_ (parental cell cycle duration). The algorithm is repeated for all cells in the population until a set number of cell divisions have been reached.

Cells without any transcripts e.g. (0,0,0) are possible in our system. They exist mainly for two reasons: 1) the low numbers of genes considered in our simplified model; 2) since parental transcripts are distributed between progeny there is probability of 2/(the total number of transcripts), that all the transcripts will end up in only one of the new cells(Zhou et al., 2011). Theoretically we have no reason to omit these cells. Second, from our perspective cells without any initial transcripts represent the most naïve theoretical state without any prior information. Early embryos such as in xenopus stages that lack zygotic transcription may be the closest to such a state (Newport and Kirschner, 1982b).

Our model was developed and simulated using Mathematica (Wolfram-Research, 2017).

### Quantification and statistical analysis

#### Gene length analysis

All protein coding genes were downloaded from Ensembl version 95 (Yates et al., 2016) using the R Biomart package version 3.5 (Durinck et al., 2009). The length of each gene was calculated using start_position and end_positions for each gene as extracted from Ensembl data.

#### Single cell analysis pipeline

Simulated datasets were pre-processed and clustered in R using the standard workflow implemented in the Seurat package. Data were log-normalized and scaled before principal component analysis (PCA) was used to reduce the dimensionality of the datasets. Due to the small number of simulated genes in the datasets, the maximum number of PCs (one fewer than the number of genes) was calculated and used in clustering. Cells were clustered using a shared nearest neighbor (SNN)-based algorithm implemented in Seurat, version 2.3.3 (Satija et al., 2015). The clustering resolution was set to 1 for all datasets, and all calculated PCs were used in the downstream clustering process. The dataset was visualized with t-SNE after clustering.

### Data and code availability

Our simulation code is available at https://github.com/BaderLab/Cell_Cycle_Theory

### KEY RESOURCES TABLE

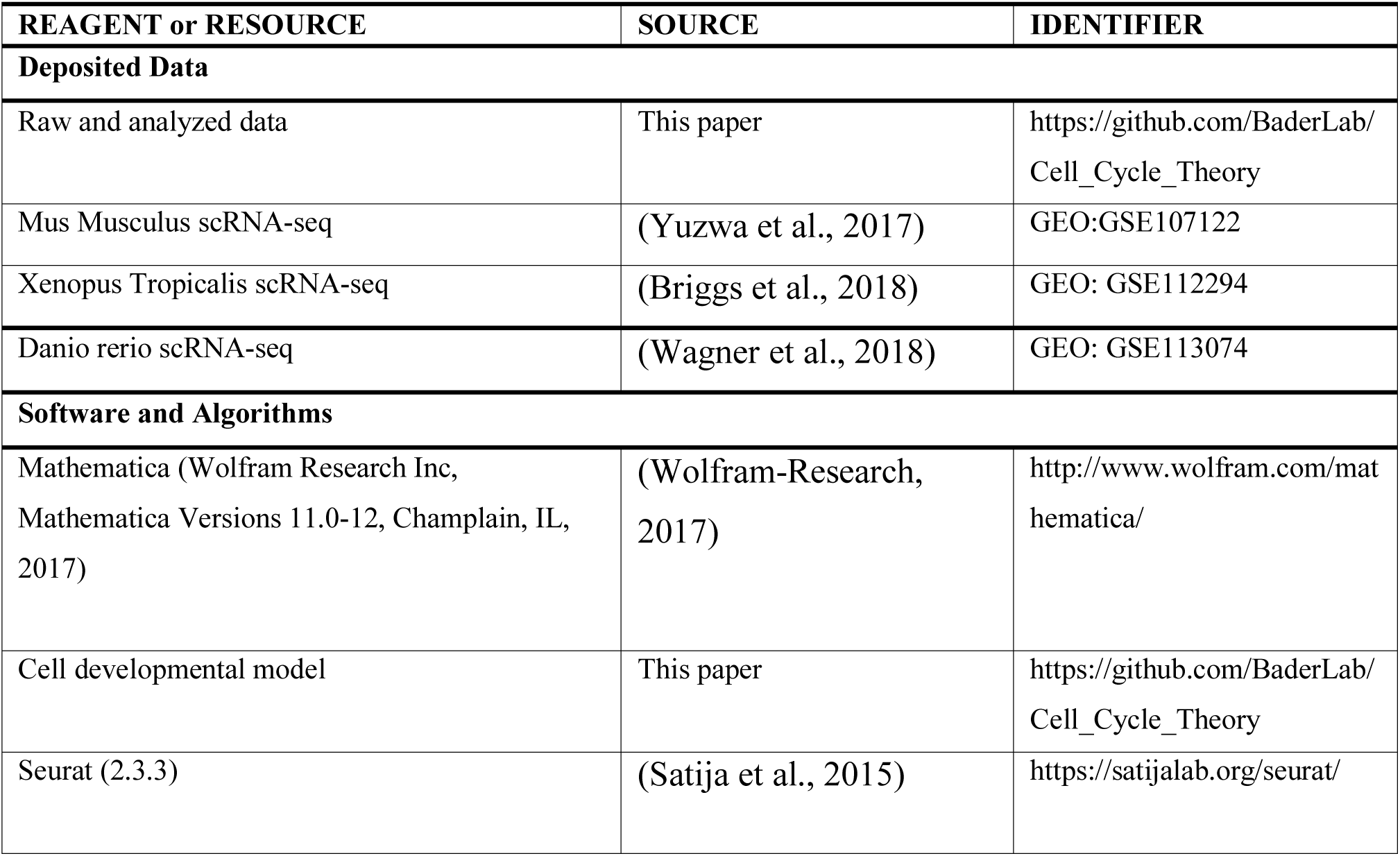
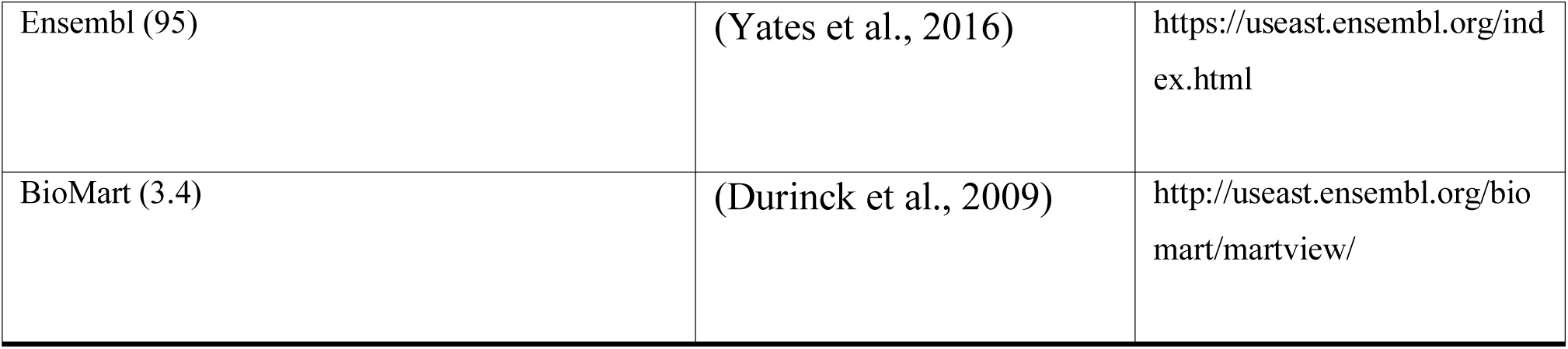

| REAGENT or RESOURCE | SOURCE | IDENTIFIER |
| --- | --- | --- |
| **Deposited Data** |  |  |
| Raw and analyzed data | This paper | https://github.com/BaderLab/Cell_Cycle_Theory |
| Mus Musculus scRNA-seq | (Yuzwa et al., 2017) | GEO:GSE107122 |
| Xenopus Tropicalis scRNA-seq | (Briggs et al., 2018) | GEO: GSE112294 |
| Danio rerio scRNA-seq | (Wagner et al., 2018) | GEO: GSE113074 |
| **Software and Algorithms** |  |  |
| Mathematica (Wolfram Research Inc, Mathematica Versions 11.0-12, Champlain, IL, 2017) | (Wolfram-Research, 2017) | http://www.wolfram.com/mathematica/ |
| Cell developmental model | This paper | https://github.com/BaderLab/Cell_Cycle_Theory |
| Seurat (2.3.3) | (Satija et al., 2015) | https://satijalab.org/seurat/ |
| Ensembl (95) | (Yates et al., 2016) | https://useast.ensembl.org/index.html |
| BioMart (3.4) | (Durinck et al., 2009) | http://useast.ensembl.org/biomart/martview/ |

## Acknowledgements

We thank Zain Patel, Brendan Innes, Derek van der Kooy, Peter Zandstra, Nika Shakiba, Janet Rossant, Eszter Posfai, Maria Shutova and Andras Nagy for thoughtful discussions about this work. This work was funded by the University of Toronto Medicine by Design initiative, by the Canada First Research Excellence Fund.

## Authors Contributions

Conceptualization, M.A.C. and G.B.; Writing-Original Draft, M.A.C. and G.B; Funding Acquisition G.B.; Formal Analysis, M.A.C., R.I. and T.T.; Methodology, M.A.C.; Investigation, M.A.C.; Visualization M.A.C., G.B. and T.T.

## Supplementary Material

**Figure S1:**
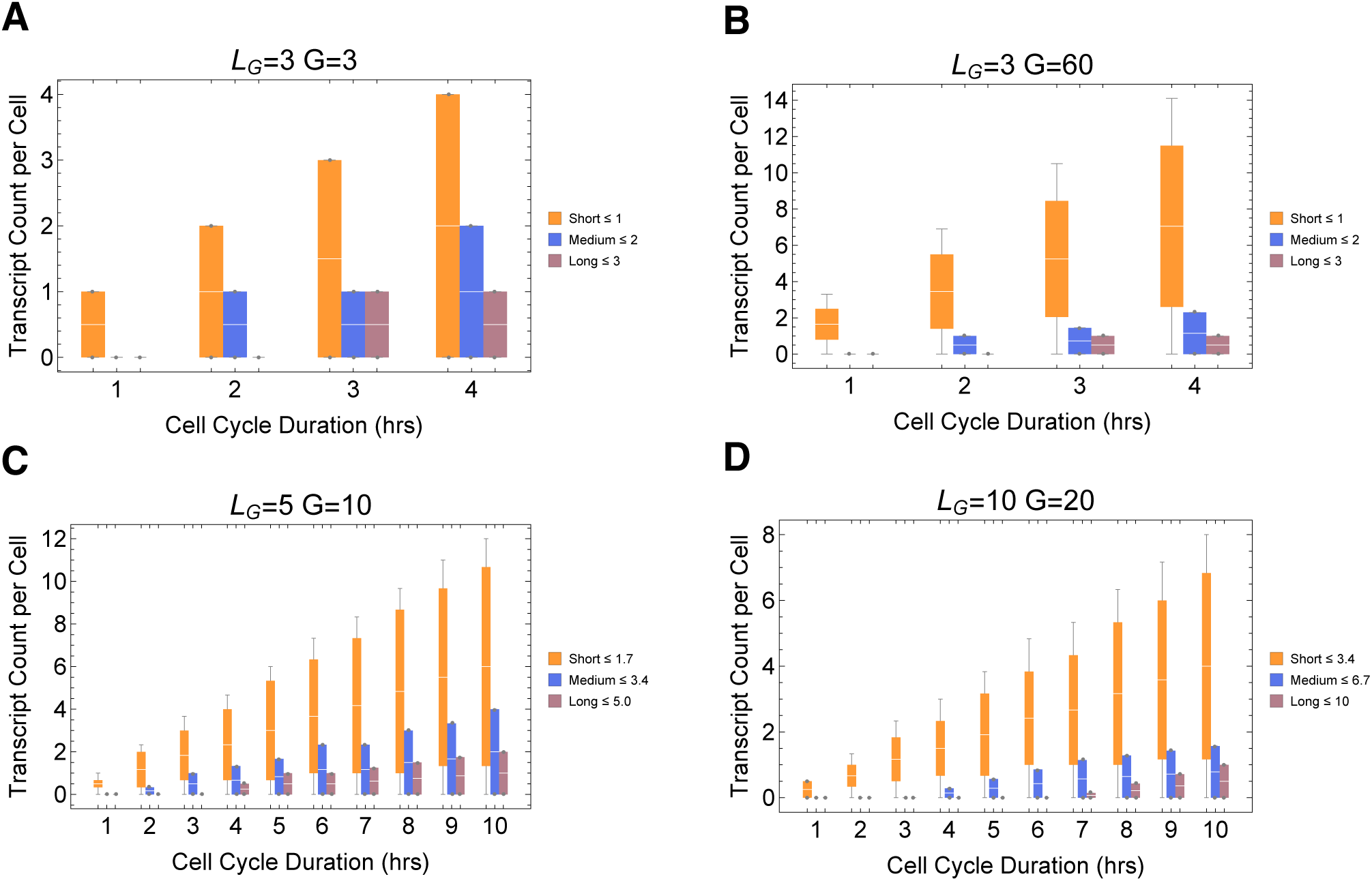
Simulations exploring the effects of cell cycle duration on transcript count per cell. We find that short genes produced more transcripts than longer genes. We conducted simulations for different gene numbers, A) G=3 B) G=60 C) G=10 D) G=20, various cell cycle durations (1-10 hrs) and maximum gene length L_G_ = (3,5 and 10). The transcriptome for each cell is subdivided into short, medium, and long genes. Each bin is averaged for each cell. Prediction from simulation shows that cells will have higher expression of short genes and longer genes irrespective the number of genes in each bin. However, simulation shows that longer cycle durations will increase relative transcript count per cell C) and D) (Other parameters ploidy=1, one cell division, Iterations= ∼5000-20000)

**Figure S 2:**
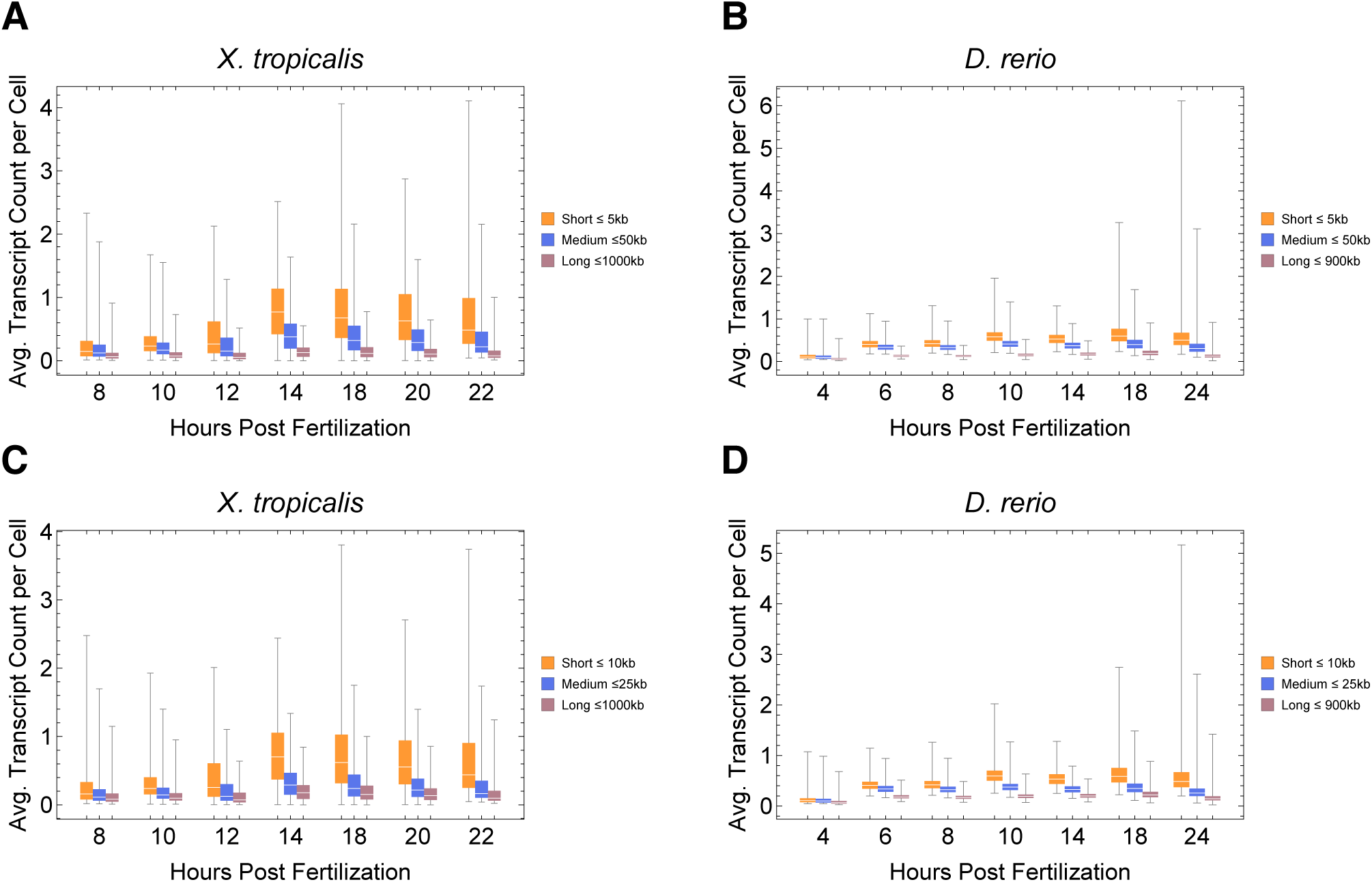
A,C) *Xenopus tropicalis* (Xenopus) single cell data obtained from GSE113074 (Lib2) and B,D) *Danio rerio* (zebrafish) single cell data obtained from GSE112294 displaying predicted patterns where short genes have a higher expression than longer genes. Genes in each cell were grouped in bins based on length and each bin was averaged per cell. A) Short genes were between 0-5 kb (1977 genes), medium genes were between 5-50 kb (7544 genes) and long genes were anything greater than 50 kb (1308 genes) B) Short genes were between 0-5 kb (3853 genes), medium genes were between 5-50 kb (12609 genes) and long genes were anything greater than 50 kb (3145 genes). Thus, the trend remains the same even if we change bin size. C) Short genes were between 0-10 kb (4174 genes), medium genes were between 5-25 kb (3607 genes) and long genes were anything greater than 25 kb (3049 genes). D) Short genes were between 0-10 kb (7595 genes), medium genes were between 5-25 kb (5899 genes) and long genes were anything greater than 25 kb (6313 genes).

**Figure S 3:**
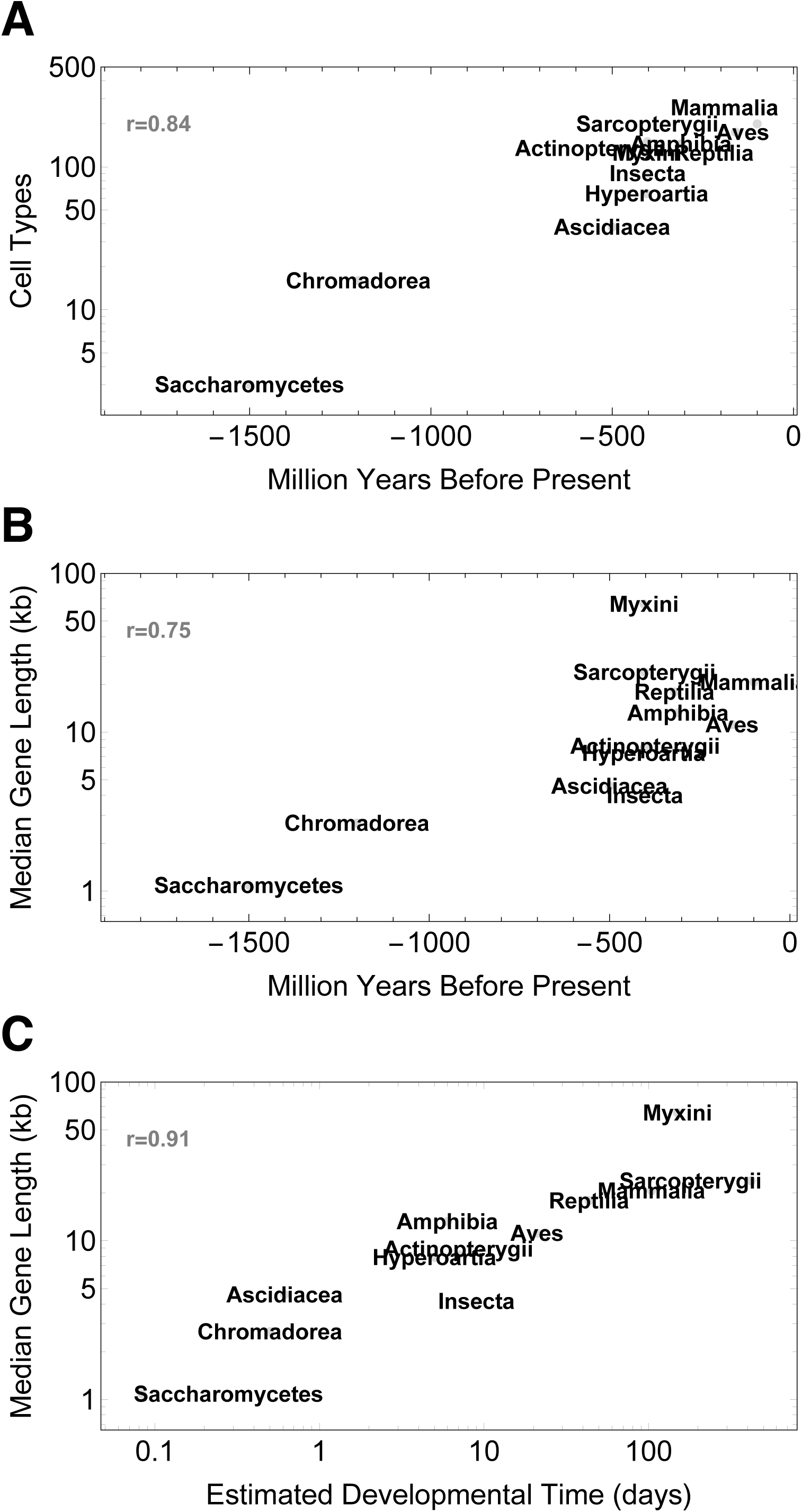
Illustrating the relationship between developmental time and median gene length across 101 species (table.xls file). For each species we calculated median gene length: All protein coding genes were downloaded from Ensembl version 95 (Yates et al., 2016) using the R Biomart package(Durinck et al., 2009). The length of each gene was calculated using start_position and end_positions for each gene as extracted from Ensembl data. Estimated developmental time was curated from encyclopedia of life or articles found in PubMed (Table S2). We used gestation time for mammals and hatching time for species who lay eggs (since it is difficult to accurately define a comparative stage for all species). We analyzed the data using a pearson correlation test shown as r.

**Figure S4:**
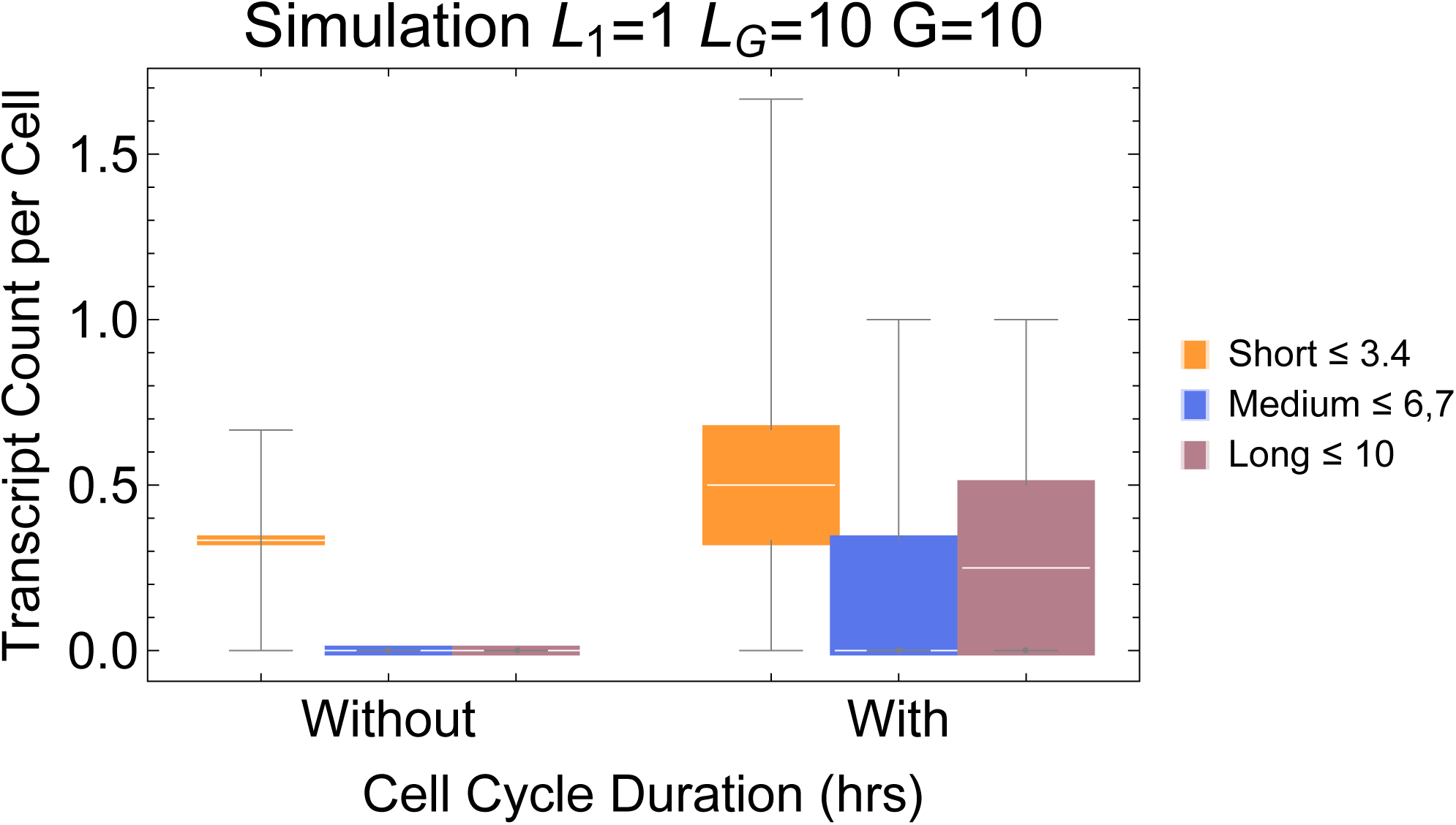
Simulation of expected transcript count when cells are initiated without maternal transcripts in comparison to cells with maternal transcripts. Parameters ploidy=1, one cell division, 1 hr cell cycle duration, iterations = 10000).

